# Mitochondrial degradation of metallothionein enables local zinc mobilization during zinc limitation

**DOI:** 10.64898/2026.06.09.731205

**Authors:** Austin K. Murchison, Julia C. Heiby, Hansen Tao, Matthew J. Matrongolo, Alessandro Ori, Monther Abu-Remaileh

## Abstract

Zinc is an essential structural and enzymatic cofactor for roughly 10% of proteins, including transcription factors, metabolic enzymes, and cytoskeletal components. It also supports critical functions across organelles such as gene regulation in the nucleus, protein folding in the endoplasmic reticulum, and energy production and antioxidant defense in mitochondria. Despite these indispensable roles, the cellular mechanism that recycles zinc to maintain homeostasis during zinc deficiency remains poorly understood. Here, we identify a biphasic response to zinc limitation, which involves the rapid degradation of the zinc-storing metallothionein followed by the degradation, in an autophagy-dependent manner, of other zinc-binding proteins. We show that metallothionein is rapidly imported into the mitochondria to be degraded by the mitoprotease LONP1. Zinc starvation leads to severe mitochondrial dysfunction and metallothionein degradation allows local zinc release to alleviate nutrient stress. Our results reveal a non-canonical, mitochondria-mediated degradation pathway for a nutrient-storing protein that mobilizes zinc locally to maintain metabolic homeostasis and establish mitochondria as active hubs for nutrient recycling.

## Introduction

Zinc is an essential metal whose abundance in biological systems. Zinc serves as a structural and catalytic cofactor for a significant number of proteins^1^. Zinc-binding proteins function in a multitude of homeostatic processes including but not limited to gene regulation, cellular metabolism and antioxidant function^2^. Around 3000 proteins in the body bind to zinc accounting for 10% of the human proteome^3^. Most intracellular zinc is protein-bound with total cellular zinc concentrations estimated at 200 to 300 DM^4^. Zinc homeostasis is maintained through a network of zinc transporters and zinc binding proteins. Import and export are tightly regulated by specific solute carrier families SLC30 (ZnT) and SLC39 (ZIP)^5^. SLC30 (ZnT) transporters regulate the flux of zinc from the cytosol into vesicles, organelles or outside of the cell. While SLC39 (ZIP) transporters are responsible for moving zinc into the cytosol from outside the cell or from organelles. Because zinc cannot be synthesized de novo, cells are dependent on environmental uptake and intracellular recycling mechanisms to maintain homeostasis during fluctuations in zinc status. Metallothionein, a small cysteine rich protein, is responsible for storage and release of intracellular zinc^6^.

Mitochondria are the primary site of adenosine triphosphate (ATP) synthesis and a central hub of cellular metabolism^7^. Zinc is indispensable for the function of mitochondria and broader cellular metabolism. Mitochondrial zinc supports oxidative phosphorylation, TCA cycle and mitochondrial respiration^8^. Heme and iron-sulfur clusters are essential cofactors of the electron transport chain in the mitochondria, and their biosynthesis requires several zinc-dependent proteins^9^. In addition, the mitochondrial metalloproteases YME1L1, AFG3L2, and OMA1 are zinc-dependent and maintain proteostasis in the mitochondria by degrading a range of substrates in the inner membrane and matrix^10^. Consequently, zinc deficiency can disrupt these proteolytic pathways and other zinc-dependent activities leading to loss of mitochondrial proteostasis and broader mitochondrial dysfunction^11–13^.

Zinc deficiency is a global nutritional health problem affecting 18% of the world’s population^14^. Zinc deficiency leads to growth stunting, immune cell dysfunction, alterations in brain function, and dysregulation of cell metabolism^15^. Furthermore, zinc deficiency has been shown to induce autophagy, a conserved mechanism to degrade macromolecules and organelles in response to stress, in Saccharomyces cerevisiae and human cells^16–18^. Proteomics results revealed many zinc proteins were degraded in zinc-deficient Saccharomyces cerevisiae. Recent studies have identified ZNG1, a novel zinc chaperone, as essential for cell survival under zinc-limited conditions^20,21^. Metallothionein levels have also been shown to change in response to zinc excess and zinc depleted conditions^22^.

Here, we show, in mammalian cells, that zinc deficiency induces a biphasic response, which includes autophagy-independent degradation of metallothionein in the early phase followed by autophagy-dependent degradation of other zinc-binding proteins. Interestingly, metallothionein is shuttled and degraded in the mitochondria by LONP1 where zinc is released to support mitochondrial metabolism. Our work identifies a potentially general process to sustain organellar nutrient demands through previously unidentified local degradation of nutrient-storing proteins.

## Results

### Zinc limitation induces autophagy-dependent global proteome remodeling

Inspired by iron recycling, which involves the selective lysosomal degradation of iron storage protein, ferritin through autophagy^23,24^, we hypothesized that zinc-recycling follows a similar pathway. To this end, we established a robust system to starve cells of zinc. We used the zinc chelators ZX1 and TPEN to deplete extracellular and intracellular zinc, respectively^25^. We performed a dose titration in wild-type PaTu-8988T cells to identify optimal concentration of each chelator by monitoring the disappearance of metallothionein, the zinc storage protein (see below) (Extended Data Fig. 1a,b). We then used quantitative proteomics to measure cellular changes in response to zinc starvation (Fig. 1a). To test autophagy dependence, we further used autophagy-deficient ATG9A-KO PaTu-8988T cells. Twenty-four hours of ZX1-induced zinc limitation caused a striking proteome remodeling in wild-type cells (Fig. 1b and Supplementary Table 1). Consistent with their preferential degradation to release zinc upon its limitation, zinc-containing proteins were significantly depleted as determined by fisher exact test (odds ratio of 2.33, p-value of 2.868 × 10^-7^) (Extended Data Fig. 1c). Autophagy-deficient cells showed an attenuated phenotype characterized by lower depletion and less overall zinc-binding proteins affected (Fig. 1c, Extended Data Fig. 1d). Of importance, zinc limitation using the intracellular zinc chelator TPEN showed similar results (Extended Data Fig. 1e,f and Supplementary Table 1).

**Figure 1.**
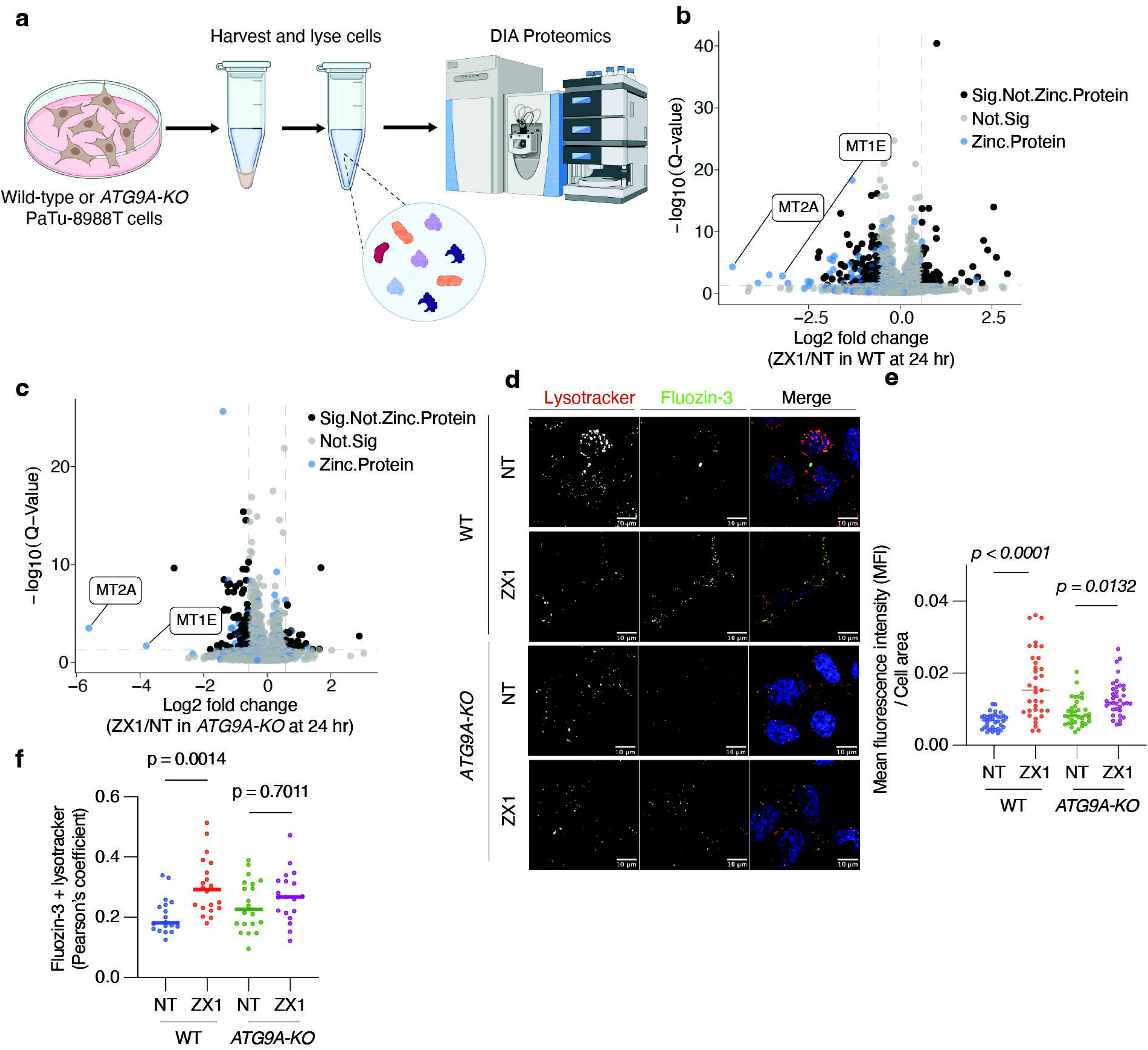
Zinc limitation induces proteome remodeling. (a) Schematic workflow of zinc starvation treatment and cell lysate collection for proteomic analysis. (b) Volcano plot of log2 fold changes (FC) in protein abundance in whole cell lysate of twenty-four hour ZX1 treated wild-type (WT) PaTu-8988T cells compared to untreated condition (n = 3 biological replicates). Dashed lines indicate absolute log2 FC > 0.58 and Q < 0.05. Blue dots highlight zinc-binding proteins, black dots show other significantly affected proteins. (c) Same as in (b) but in ATG9A-KO PaTu-8988T cells (n = 3 biological replicates). (d) Representative images from confocal microscopy using Fluozin-3 (green) and Lysotracker Red D99 (red) in wild-type and ATG9A-KO PaTu-8988T cells under ZX1 treatment. Nuclei were stained with Hoechst 33342 (blue). Wild-type and ATG9A-KO cells were treated with ZX1 (500 μM). NT indicates no ZX1 condition. Scale Bar = 10 μm. (e) Quantification of Fluozin-3. Mean fluorescence intensity (MFI) normalized by cell area in each condition (36 cells were analyzed per condition) (n = 3 biological replicates). One-way ANOVA was used to compare experimental conditions. (f) Quantification of the colocalization of labile zinc ion (Fluozin-3) and lysosomes (Lysotracker Red D99) (36 cells were analyzed per condition). (n = 3 biological replicates). Line represents mean value. One-way ANOVA was used to compare experimental conditions.

To visualize free zinc localization upon zinc limitation, we used Fluozin-3 AM, a cell-permeant Zn^2+^ dye. After twenty-four hours in zinc-limited media, labile zinc levels increased in the cell (Fig. 1d,e). This increase, however, was attenuated in autophagy-deficient cells (Fig. 1d,e). Strikingly, lysosomal zinc pools were significantly increased in wild-type cells while no similar increase is observed in autophagy-deficient cells (Fig. 1f). These results suggest an autophagy-dependent release of zinc through lysosomal degradation of zinc-containing proteins after twenty-four hours of zinc starvation.

### A biphasic cellular response to zinc limitation

To better understand the cellular response to zinc limitation, we examined the temporal profiles of proteins that were significantly altered after twenty-four hours of treatment (Fig. 2a and Supplementary Table 2). As expected, most zinc-binding proteins were gradually degraded in an autophagy-dependent manner, but only at later time points (≥ 8 hours) (Fig. 2a,b). To our surprise, zincDstoring metallothioneins disappeared earlier and more rapidly than other zincDbinding proteins (Fig. 2a,c). Moreover, unlike other zinc-binding proteins, their complete clearance from the cell was comparable in wild-type and ATG9A-KO cells (Fig. 2c).

**Figure 2.**
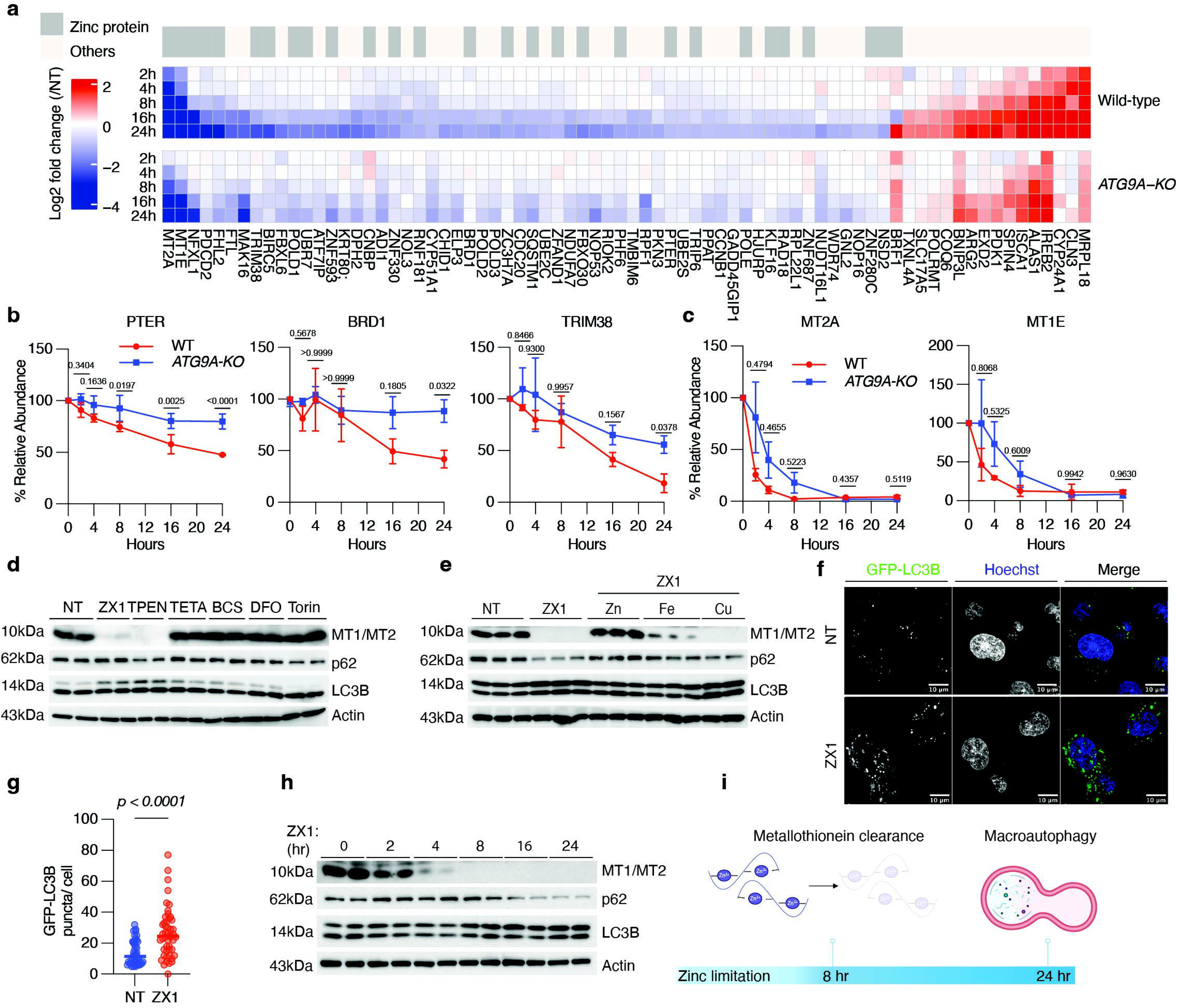
A biphasic cellular response to zinc limitation. (a) Heatmap comparing Fold Changes (FC) over a twenty-four hour time course of proteins with FC above 2 or below –2 (Q ≤ 0.05) at twenty-four hours of ZX1 treatment. Wild-type and ATGA9-KO FC are plotted separately. Zinc proteins are labeled above (n = 3 biological replicates). (b) Line plots representing zinc proteins PTER, BRD1, and TRIM38. Relative abundance is normalized to protein abundance at zero hour, wild-type is labeled in red and ATG9A-KO is labeled in blue. Two-way ANOVA was used to compare relative abundance in wild-type and ATG9A-KO at each time point (n = 3 biological replicates). Statistical significance is noted above each time point. (c) Same as in (b) for MT2A and MT1E. (d) Immunoblot analysis of lysates from PaTu-8988t cells treated with a panel of chelators. ZX1(Zn^2+^), TPEN (Zn^2+^), TETA (Cu^2+^), BCS (Cu^+^), DFO (Fe^3+^), and the mTOR inhibitor Torin1 were added to cells for eight hours. p62 and LC3B were used as autophagy markers. Antibody recognizing MT1 and MT2 was used to monitor metallothioneins and D-Actin was used as loading control. (e) Immunoblot analysis of lysates from PaTu-8988t cells treated with ZX1(500 DM) and equal molar ZnCl_2_, Ferric Ammonium Chloride (FAC), or CuSO_4_ for twenty-four hours. p62 and LC3B were used as autophagy markers. Antibody recognizing MT1 and MT2 was used to monitor metallothioneins and D-Actin was used as loading control. (f) Representative image of autophagy measurement using PaTu-8988t cells stably expressing GFP-LC3B reporter. GFP-LC3B is shown in green. Nuclei are stained with Hoechst 33342 in blue. Cells were treated with ZX1(500 μM) for twenty-four hours. NT=No treatment. (g) Quantification of GFP-LC3B puncta in (f) (44-48 cells were analyzed per condition) (n = 3 biological replicates). Line represents mean value. Unpaired t-test was used to compare experimental conditions. (h) Immunoblot analysis of twenty-four time-resolved ZX1(500 DM) treatment in PaTu-8988T cells. p62 and LC3B were used to monitor autophagy. Antibody recognizing MT1 and MT2 was used to monitor metallothioneins and D-Actin was used as loading control. (i) Two-phase response to zinc limitation involves the rapid degradation of zinc-storing metallothioneins within an eight-hour time window followed by the induction of autophagy after the depletion of metallothionein proteins.

Next, we sought to confirm this degradation was specific to decreased zinc levels. We leveraged an antibody that recognizes both MT1A and MT2A proteins (Extended Data Fig. 2a) to test the impact of known copper and iron chelators including BCS (Cu^+^), TETA (Cu^+^/Cu^2+^) and Deferoxamine (Fe^3+^) along with the mTOR inhibitor Torin1. ZX1 and TPEN showed robust clearance of metallothionein while depletion of copper and iron had no effect on metallothionein levels (Fig. 2d). Similarly, Torin1 showed no effect on metallothionein, thus ruling out metallothionein clearance as a nonspecific response to mTOR inhibition and induction of autophagy (Fig. 2d). To test the selectivity of the ZX1 chelator, we co-treated the ZX1-treated cells with equal molar zinc (II) chloride, ferrous (II) chloride, and copper sulfate. Co-treatment with equal molar zinc fully rescued metallothionein clearance (Fig. 2e). Co-treatment with equal molar iron slightly rescued metallothionein degradation while co-treatment with copper had no effect (Fig. 2e).

Our data showed a robust induction of autophagy in response to zinc limitation for twenty-four hours as assessed by lipidated LC3B and p62 levels (Fig. 2d,e), which we also confirmed using a GFP-LC3B reporter as an orthogonal approach (Fig. 2f,g).

To understand the link between metallothionein clearance and autophagy induction, we performed a time-resolved experiment that confirmed degradation of metallothionein and autophagy induction were occurring at two distinct times following zinc starvation. Metallothionein is cleared within the first eight hours followed by the induction of autophagy (Fig. 2h). These data are consistent with our proteomic analysis showing major effects on zinc-binding proteins, except for metallothioneins, in an autophagy-dependent manner at this later stage (Fig. 2a-c). Altogether, these results suggest a biphasic response to zinc starvation, which starts with rapid clearance of metallothionein in the “early phase”, followed by the induction of autophagy to degrade other zinc-binding proteins in the “late phase” (Fig. 2i).

### Metallothionein clearance is independent of autophagy, lysosome, and proteasome

To validate our observations from the proteomics experiment (Fig. 2a,c), we monitored metallothionein clearance in both ATG9A– and ATG7-KO cells using immunoblotting. While a minor slowdown was observed in ATG9A-KO cells, metallothionein degradation was autophagy independent within the first eight hours of zinc starvation (Fig. 3a,b). Proteins can be degraded by the lysosome through mechanisms independent of canonical autophagy such as chaperone-mediated autophagy or by direct import into the lysosome^26,27^. However, inhibiting lysosomal function using Bafilomycin A1 and Chloroquine that raise lysosomal pH only slightly attenuated metallothionein clearance at four hours, which then completely disappeared within the eight hours window (Fig. 3c). Next, we tested if the Ubiquitin-Proteasome System (UPS) is responsible for the clearance using two well-established proteasome inhibitors bortezomib and MG-132^28–30^. Both potently inhibited proteasome function as indicated by the accumulation of ubiquitinylated proteins but failed to prevent metallothionein clearance (Fig. 3d). Altogether, these data show that canonical protein degradation pathways are dispensable for metallothionein clearance.

**Figure 3.**
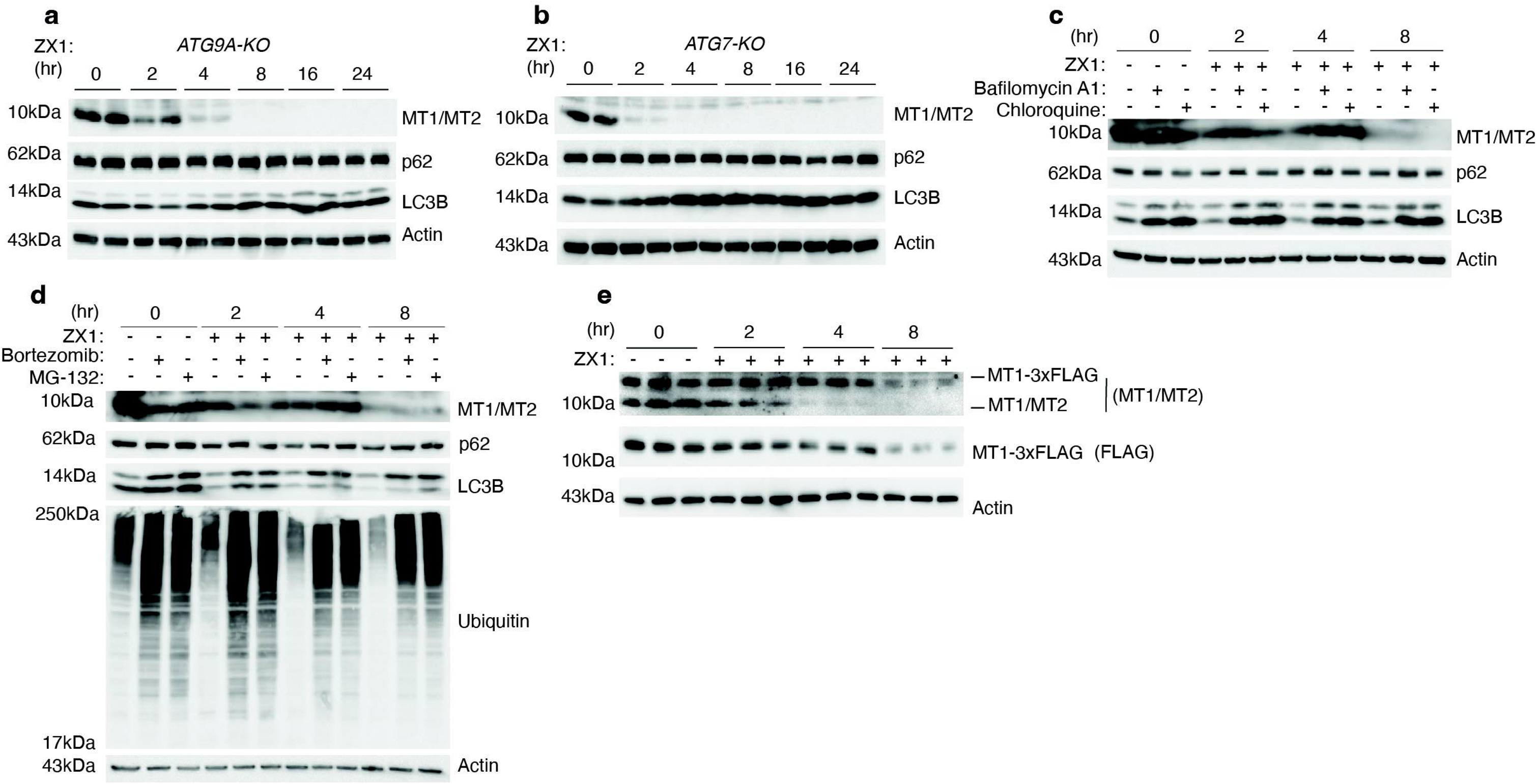
Metallothionein clearance is independent of autophagy, lysosome, and proteasome. (a) Immunoblot analysis of twenty-four time-resolved treatment with ZX1 (500 DM) in ATG9A-KO PaTu-8988T cells. p62 and LC3B were used to monitor autophagy. Antibody recognizing MT1 and MT2 was used to monitor metallothioneins and D-Actin was used as loading control. (b) Immunoblot analysis of twenty-four time-resolved treatments with ZX1 (500 DM) in ATG7-KO PaTu-8988T cells. p62 and LC3B were used to monitor autophagy. Antibody recognizing MT1 and MT2 was used to monitor metallothioneins and D-Actin was used as loading control. (c) Immunoblot analysis of time-resolved zinc depletion with ZX1 (500 DM) in wild-type PaTu-8988Ts cotreated with lysosome inhibitors Bafilomycin A1 (500 nM) and Chloroquine (50 DM). p62 and LC3B were used to monitor autophagy. Antibody recognizing MT1 and MT2 was used to monitor metallothioneins and D-Actin was used as loading control. (d) Immunoblot analysis of time-resolved zinc depletion with ZX1 (500 DM) in wild-type PaTu-8988Ts cotreated with proteasome inhibitors Bortezomib (500 nM) and MG-132 (5 DM). Ubiquitin antibody staining was used to confirm proteasome inhibition. p62 and LC3B were used to monitor autophagy. Antibody recognizing MT1 and MT2 was used to monitor metallothioneins and D-Actin was used as loading control. (e) Immunoblot analysis of time-resolved zinc depletion with ZX1 (500 DM) in PaTu-8988T cells expressing MT1-3xFLAG reporter. Antibody recognizing MT1 and MT2, as well as MT1-3xFLAG was used. MT1A-3xFLAG was also monitored using anti-Flag antibody and D-Actin was used as loading control.

Finally, we sought to test if metallothionein clearance is occurring at the protein level rather than a result of transcriptional inhibition. Therefore, we constructed a MT1-3xFLAG reporter. Similar to the endogenous metallothionein, the reporter was completely cleared in response to ZX1 and TPEN treatment at twenty-four hours (Extended Data Fig. 3a), and a time-resolved experiment showed a significant degradation by eight hours despite overexpression (Fig. 3e). To rule out intracellular degradation entirely, we used a commercially available protease cocktail containing pepstatin A, leupeptin, E-64, aprotinin, and bestatin to target all intracellular proteases^31^. Surprisingly, this treatment significantly inhibited the degradation of the metallothionein reporter and spared a significant fraction of the endogenous one at eight hours (Extended Data Fig. 3b). These results support a possible novel degradation pathway that is responsible for metallothionein clearance and zinc release under starvation.

### Metallothionein is imported into the mitochondria upon zinc limitation for degradation by LONP1

To identify the molecular machinery necessary to facilitate metallothionein clearance, we used our MT1-3xFLAG reporter for co-immunoprecipitation (co-IP) coupled with proteomics to identify molecular interactors (Fig. 4a). We validated the enrichment of the reporter using anti-Flag magnetic beads (Extended Data Fig. 4a). Proteomic analysis revealed TOMM70, a major component of the TOM complex which is responsible for importing proteins synthesized in the cytosol into the mitochondria, as an interactor (Fig. 4b and Supplementary Table 3)^32^. Furthermore, we found that among the differentially enriched protein interactors at four hours of zinc starvation is LONP1 (Fig. 4c), a mitochondrial protease localized to the mitochondrial matrix^33,34^. Based on these findings, we hypothesized that metallothionein is recruited to the mitochondria for degradation by LONP1. First, we tested if mitochondrial import is required for metallothionein clearance. To this end, we inhibited mitochondrial protein import in zinc-limited cells using FCCP, a mitochondrial uncoupler that disrupts the TOM complex by ablation of the membrane potential^35,36^. Indeed, FCCP treatment completely inhibited metallothionein degradation (Fig. 4d). Furthermore, metallothionein degradation was also inhibited by mitoblock-6, a mitochondrial import inhibitor targeting the Erv1-Mia40 pathway^37^ (Extended Data Fig. 4b).

**Figure 4.**
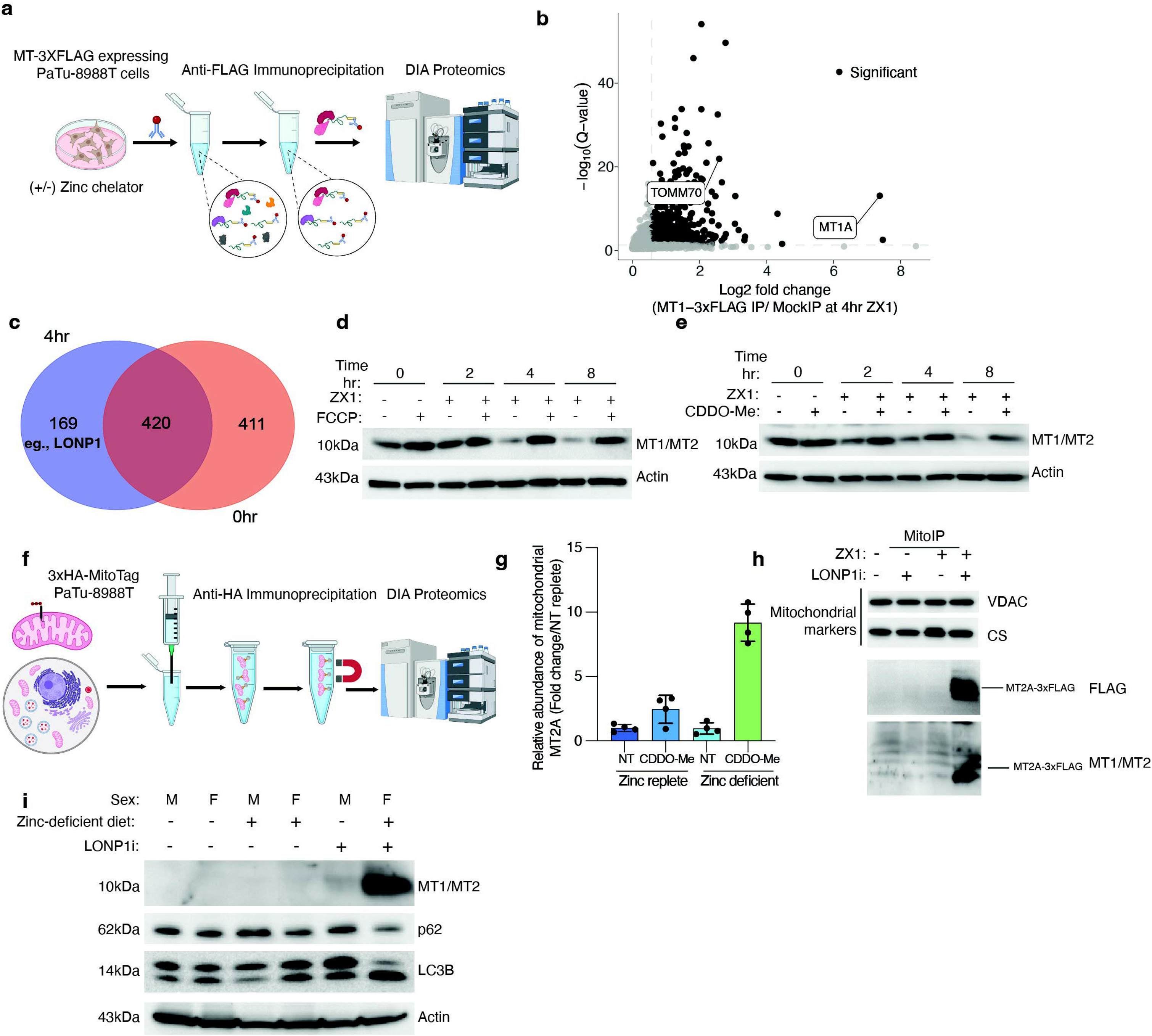
Metallothionein is imported into the mitochondria upon zinc limitation for degradation by LONP1. (a) Representative workflow for anti-Flag co-immunoprecipitation (CoIP) from PaTu-8988T cells stably expressing MT1-3xFLAG. Wild-type PaTu-8988T cells not expressing metallothionein reporter were used as background control. (b) Volcano plot of log2 fold changes (FC) in protein abundance between CoIP from reporter expressing cells and from those that do not express it shows potential protein interaction partners of MT1-3xFLAG after four-hour ZX1(500 DM) treatment. (c) Venn Diagram comparing interaction partners at four hours of ZX1(500 DM) compared to those enriched in zinc replete control (zero hours) (all show log2FC>0.58 and Q <= 0.05). Proteases found to be interactors at four hours are listed. (d) Immunoblot analysis of wild-type PaTu-8988T cells treated with ZX1(500 DM) at each indicated time and co-treated with mitochondria uncoupler FCCP (10 DM). (e) Immunoblot of wild-type PaTu-8988T cells treated with ZX1(500 DM) at each indicated time and co-treated LONP1 protease inhibitor CDDO-Me (3 DM). (f) Workflow of MitoIP coupled with proteomics. (g) Relative abundance of MT2A in the mitochondrial fraction determined by proteomics from cells growing in zinc replete and four hour zinc-limited media co-treated with LONP1 inhibitor CDDO-Me (3 DM). (h) Immunoblot analysis of MitoIP from PaTu-8988T cells expressing MT2A-3xFLAG and growing in zinc replete or four-hour zinc-limited media and co-treated with LONP1 inhibitor CDDO-Me (3 DM). (i) Immunoblot analysis of livers from wild-type mice treated for three days with CDDO-Me before switching to zinc-deficient diet for six days. Mice were IP injected daily with 2.5 mg/kg CDDO-Me in DMSO for a total of nine days. p62 and LC3B were used to monitor autophagy. Antibody recognizing MT1 and MT2 was used to monitor metallothioneins and D-Actin was used as loading control.

To test if LONP1 is responsible for degrading metallothionein, we used CDDO-Me, a well-established LONP1 inhibitor^38^. Indeed, CDDO-Me inhibited metallothionein degradation under zinc starvation (Fig. 4e). LONP1 is a highly essential gene^39^, thus generating single cell KO clones is extremely challenging. However, population level CRISPR KO cells showed significantly slower degradation kinetics compared to wild-type cells despite incomplete loss of LONP1 (Extended Data Fig. 4c). Altogether, these data suggest that metallothionein is imported into mitochondria for degradation, potentially by LONP1. To validate these findings, we utilized the MitoIP technique, which allows rapid isolation of mitochondria using a genetically encoded MitoTag (Fig. 4f)^40^. Consistent with our model, we observed a striking enrichment of the endogenous MT2A in the mitochondria when LONP1 is inhibited (Fig. 4g, Extended Data Fig. 4d, and Supplementary Table 4). Similarly, we were able to capture strong enrichment of our metallothionein reporter in the mitochondria upon zinc starvation and LONP1 inhibition (Fig. 4h). These results confirm that metallothionein is imported into the mitochondria to be rapidly degraded by LONP1.

Finally, to test if mitochondria-mediated degradation of metallothionein by LONP1 is relevant in organismal homeostasis, we sought to validate our observations in vivo. To this end, mice were treated with an intraperitoneal injection of LONP1 inhibitor and fed zinc-deficient chow for six days. Consistent with our in vitro data, LONP1 inhibition in zinc-deficient diet fed mice caused a strong increase in cellular metallothionein, indicative of dysfunctional degradation by LONP1 (Fig. 4i). This result demonstrates mitochondria-mediated degradation of metallothionein is relevant for organismal adaptation to zinc deficiency.

### Metallothionein degradation rapidly releases zinc in the mitochondria during zinc limitation

Zinc supports vital mitochondrial processes and is essential for mitochondrial homeostasis^41,42^. Consistently, we found that zinc starvation leads to mitochondria fragmentation (Fig. 5a,b). Additionally, mitochondrial respiration was severely decreased under zinc-limited conditions, with oxygen consumption rates reaching background levels after twenty-four hours of ZX1 treatment (Fig. 5c,d). Based on these observations, we hypothesized that metallothionein is recruited to mitochondria to locally provide zinc to support metabolic demands. To this end, we used the zinc-specific probe Zinpyr-1 to visualize labile pools of zinc in the cell. While in zinc-replete conditions only a small fraction of zinc overlapped with Mitotracker red, a mitochondria-specific dye, extracellular zinc limitation leads to significant increase in free mitochondrial zinc (Fig. 5e,f). To test whether blocking metallothionein import in the mitochondria or its mitochondrial degradation inhibit zinc release, we repeated the assay using MitoBright, a mitochondrial dye that is insensitive to membrane potential, to avoid treatment-dependent changes in mitochondrial labeling. Indeed, zinc limitation increased the labile pools of zinc in the mitochondria while inhibiting mitochondrial potential or LONP1 using FCCP and CDDO-Me, respectively, significantly decreased mitochondrial zinc release (Fig. 5g,h). These results strongly demonstrate that metallothionein is targeted to mitochondria upon zinc limitation to locally release stored zinc to support mitochondrial metabolism.

**Figure 5.**
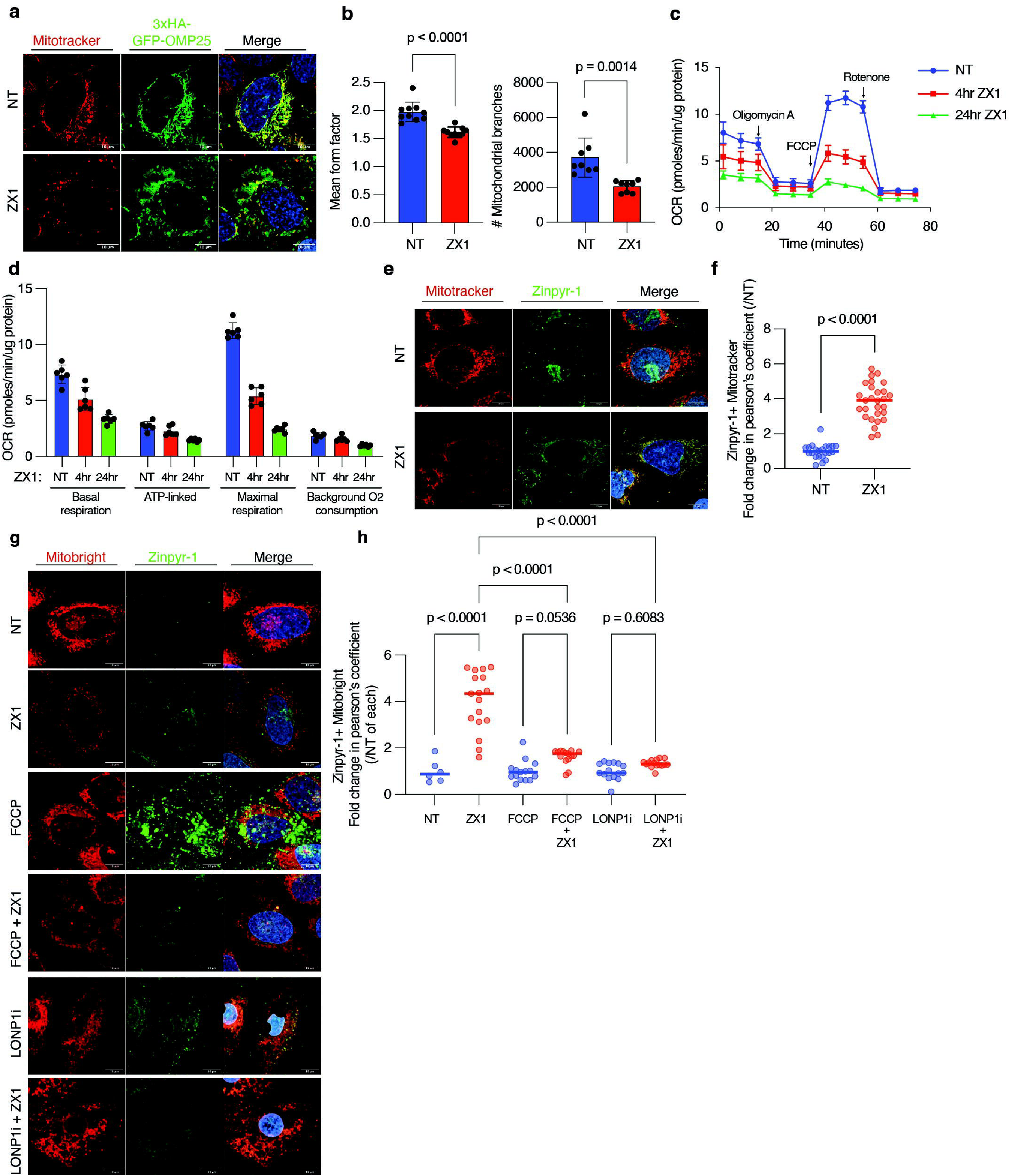
Metallothionein degradation rapidly releases zinc in the mitochondria during zinc limitation. (a) Representative image showing mitochondria morphology in U2OS cells stably expressing 3xHA-GFP-OMP25. Mitochondria were also labeled with Mitotracker red. Nuclei are stained with Hoescht 33342 in blue. Cells were treated with ZX1(500 DM). (b) Quantification of mitochondria morphology using U2OS cells from (a). 10 fields were used (n = 2 biological replicates). NT=No treatment. ZX1= ZX1(500 DM) for twenty-four hours. Mean form factor: Quantification of mitochondrial elongation and shape complexity. Number of branches: Measurement of mitochondrial network connectivity and fusion. (c) Mitochondrial oxygen consumption rates (OCR) in PaTu-8988Ts treated with ZX1. No Treatment (n = 6), four-hour ZX1 (n = 6), and twenty-four ZX1 (n = 6). OCR reads were normalized to total protein (DM). Results represent mean ± SEM. (d) Bar plot representation of OCR in wild-type PaTu-8988Ts from (c). Results represent mean ± SEM. (e) Representative images of U2OS cells stained with zinc-specific dye Zinpyr-1 (5 DM), Mitotracker Red (100 nM) and Hoechst 33342 (blue). U2OS were zinc replete (NT) or treated with ZX1 for four-hours to visualize zinc release in the mitochondria. Scale Bar = 10 Dm. (f) Quantification of colocalization (Pearson’s coefficient) of Zinpyr-1(zinc pools) and mitotracker red (mitochondria) per field (n = 20 fields from n = 3 biological replicates) from (e). Data is represented as fold change. T-test was used to compare colocalization of Zinpyr-1 and Mitotracker between ZX1-treated and untreated conditions. (g) Representative images of U2OS cells stained with Zinpyr-1 (5DM), Mitobright Deep red (100 nM) and Hoechst 33342 (blue). U2OS were zinc replete (NT) or treated with ZX1 for four hours to visualize zinc release in the mitochondria. Scale Bar = 10 Dm. (h) Quantification of colocalization (Pearson’s coefficient) of Zinpyr-1 (5 DM) and Mitobright Deep red (100 nM) per field (n = 6-17 fields from n = 3 biological replicates) from (g). Data is represented as fold change between ZX1 treatment and the no ZX1 control in each group. One-way ANOVA was used to compare colocalization of Zinpyr-1 and Mitobright deep red in noted conditions.

## Discussion

Our work identifies a novel biphasic response to zinc limitation in the mammalian system. In an early phase, metallothionein is rapidly imported into the mitochondria and degraded by LONP1 to locally release zinc. In a later phase, zinc-binding proteins are more broadly degraded through the autophagy-lysosome axis. These findings suggest that cells engage a temporally ordered zinc-sparing program in which mitochondrial zinc needs are prioritized before bulk recycling pathways are activated.

This model expands the classical view that nutrient recycling is centered in the lysosome^43–45^. While lysosomal autophagy contributes to the degradation of zinc-binding proteins during prolonged zinc limitation, our data show that mitochondria can directly degrade the zinc-storing protein metallothionein to rapidly generate a local pool of labile zinc. This local release may provide a fast mechanism to preserve mitochondrial function under nutrient stress, particularly because zinc-dependent mitochondrial processes are acutely vulnerable.

Several observations are consistent with this idea. Zinc limitation causes severe mitochondrial dysfunction, while metallothionein is recruited to mitochondria, degraded in a LONP1-dependent manner, and accompanied by increased mitochondrial labile zinc. These findings suggest that local zinc release may help sustain essential mitochondrial pathways, potentially including heme biosynthesis and iron-sulfur cluster-dependent respiration^8,46–48^, although defining the specific zinc-dependent processes supported by this pathway will require further study.

More broadly, mitochondrial degradation of metallothionein raises the possibility that organelles can locally mobilize stored nutrients through degradation of small storage or chaperone proteins. It will be important to determine whether analogous mechanisms exist for other metals and whether similar pathways operate for mitochondrial ferritin^49,50^ or copper-binding proteins^51–53^ under defined metabolic stresses.

Finally, an important unresolved question that emerges from our work is how zinc limitation is sensed and how that signal triggers metallothionein recruitment to mitochondria. Determining whether the initiating signal arises in the cytosol, within mitochondria, or through inter-organelle communication will be important for understanding how cells prioritize zinc allocation during nutrient stress.

Altogether, this study shows that mitochondria are not only consumers of recycled nutrients but can directly participate in nutrient mobilization by degrading a storage protein locally in order to protect organelle metabolism during nutrient stress.

## Methods and Materials

### Cell culture

All cell lines and their derivatives were cultured in Dulbecco’s Modified Eagle Medium (DMEM; Gibco) supplemented with 10% heat inactivated fetal bovine serum (FBS) (Thermo Fisher Scientific), GlutaMAX (Gibco), and penicillin and streptomycin (Thermo Fisher Scientific). For zinc-limited culture media, ZX1 (Strem Chemicals) and TPEN (Cayman Chemicals) were added to DMEM with 10% FBS, GlutaMAX, and penicillin and streptomycin. All cell lines were maintained at 37°C and 5% CO_2._ All cell lines were frequently tested for mycoplasma.

### Plasmid Construction

Gene blocks encoding MT1A and MT2A with a C-terminal 3xFLAG epitope tag were ordered from Twist Biosciences. pMX-3xFLAG-eGFP-OMP25 (Addgene #83354) was used to clone pMX-MT1A-3xFLAG retrovirus plasmid. MT1A gene block was cloned into pMXs-IRES-Blast-3xFLAG-GFP vector using BamHI and NotI restriction cloning. pLJM1-Empty (Addgene #91980) was used to clone pLJM1-MT2A-3xFLAG lentivirus plasmid. MT2A gene block was cloned into pLJM1-Empty vector using AgeI and EcoRI restriction cloning.

### Recombinant virus production and transduction

Viral HEK293T cells were grown to 60% confluency and transfected with appropriate plasmids to generate lentivirus and retrovirus. To generate transgene lentivirus, lentivirus constructs were transfected with psPAX2 and pCMV-VSV-G. Retrovirus constructs were transfected with pCMV-VSV-G and packaging plasmid. Viral plasmids were transfected in viral HEK293Ts using XtremeGene 9 (Roche). Sixteen hours after transfection, culture medium was replaced with DMEM supplemented with 30% FBS and penicillin/streptomycin. The virus-containing supernatant was harvested after 48 hours and centrifuged to remove cell debris.

Two million cells were plated in six-well plates in DMEM with the addition of polybrene (8 Dg/ml) and 100 to 250 Dl of virus-containing medium. Spin infection was performed at 2200 rpm for 45 min at 37°C, and cells were incubated with virus for 16 hours before adding fresh culture medium containing the relevant antibiotic for selection for at least 72 hours.

### Western Blot

Cell lysates were resolved by SDS-polyacrylamide gel electrophoresis at 120 V and transferred to polyvinylidene difluoride membranes for 2 hours at 40 V. Membranes were blocked with 5% BSA (bovine serum albumin) in TBST buffer (tris-buffered saline with Tween 20) for 30 min and then incubated overnight with primary antibodies in 5% BSA in TBST at 4°C. After incubation, membranes were washed three times with TBST for 30 min for wash 1 and 5 min for wash 2 and 3. The membranes were incubated with the appropriate horseradish peroxidase secondary antibodies diluted in 1:1000 in 5% BSA in TBST for 1 hour at room temperature. Membranes were washed three times with TBST and visualized using ECL2 Western Blotting substrate on a ChemiDoc MP imaging system.

| Antibodies used for western blot |  |  |  |
| --- | --- | --- | --- |
| Antibody | Vendor | Cat# | Dilution |
| Metallothionein | Novus Biologics | NBP1-97493 | 1:500 |
| β-Actin | Santa Cruz<br>Biotechnology | SC-47778 | 1:10000 |
| LC3B | Cell Signaling | 2775S | 1:1000 |
| SQSTM1/p62 | Cell Signaling | 5114S | 1:1000 |
| ATG9A | Cell Signaling | 13509S | 1:1000 |
| ATG7 | Cell Signaling | 8558S | 1:1000 |
| Ubiquitin | Cell Signaling | 3936S | 1:1000 |
| DYKDDDDK (Flag)<br>tag | Cell Signaling | 14793S | 1:500 |
| LONP1 | Proteintech | 15440-1-AP | 1:1000 |
| Anti-Mouse-HRP | Cell Signaling | 7076 | 1:1000 |
| Anti-Rabbit HRP | Cell Signaling | 7077 | 1:1000 |

### Mitochondrial isolation

Mitochondria purification was done as previously described with slight modification^40^. Briefly, at the time of harvest, cells were washed twice with ice-cold KPBS containing protein and phosphatase inhibitors. Cells were scraped into 1 ml of ice-cold KPBS with protein and phosphatase inhibitors. Cells were pelleted by centrifugation at 1000 xg for 2 min at 4°C. Following centrifugation, cells were resuspended in 500 Dl of fresh KPBS and 12.5 Dl of the cell suspension was taken for whole-cell fraction analysis. The remaining cell suspension was lysed by passing through a 29.5-gauge insulin syringe, then diluted with additional 450 Dl and centrifuged at 1000 xg for 2 min at 4°C to pellet cell debris. The supernatant containing mitochondria was transferred to 100 Dl of Pierce anti-HA magnetic beads and mixed for 5 min at 4°C. The beads bound mitochondria were washed 3 times with ice-cold KPBS and extracted in Triton lysis buffer (1% Triton X-100, 40 mM sodium chloride, 2 mM EDTA. 1.5 mM sodium orthovanadate, 30 mM sodium fluoride, 10 mM sodium pyrophosphate, 10 mM sodium beta-glycerophosphate).

### Sample preparation for proteomics analysis

For proteomics analysis, eluates from Co-IPs, MitoIPs, as well as aliquots of the matched starting lysates were thawed on ice and resuspended in 3x lysis buffer (100 mM HEPES, 50 mM DTT and 5% SDS final concentration). Samples were then sonicated (Bioruptor Plus, Diagenode, Belgium) for 10 cycles (30 s on, 60 s off) at high setting, at 20°C, followed by boiling at 95°C for 7 min. Reduction was followed by alkylation with iodoacetamide (IAA, final concentration 15 mM) for 30 min at room temperature in the dark. Samples were acidified with phosphoric acid (final concentration 1%), and six times the sample volume of S-trap binding buffer was added (100 mM Triethylammonium biscarbonate buffer, TEAB, and 90% methanol). Samples were bound on 96-well S-trap micro plate (Protifi) and washed three times with binding buffer. Trypsin in 50 mM TEAB pH 8.5 was added to the samples (1 μg per sample) and incubated for 1 h at 47°C. The samples were eluted in three steps with 50 mM TEAB pH 8.5, elution buffer 1 (0.2% formic acid in water) and elution buffer 2 (50% acetonitrile and 0.2% formic acid). The eluates were dried using a speed vacuum centrifuge (Eppendorf Concentrator Plus, Eppendorf AG, Germany) and stored at –20°C. The peptides were reconstituted in 5% (v/v) acetonitrile, 0.1% (v/v) formic acid to a peptide concentration of approximately 1 Dg/Dl and spiked with iRT peptides (Biognosys, 1:2 ratio). The samples were loaded on Evotips (Evosep) according to the manufacturer’s instructions. In short, Evotips were first washed with Evosep buffer B (acetonitrile, 0.1% formic acid), conditioned with 100% isopropanol and equilibrated with Evosep buffer A. Afterwards, the samples were loaded on the Evotips and washed with Evosep buffer A. The loaded Evotips were topped up with buffer A and stored until the measurement.

### LC-MS Data independent analysis (DIA)

Peptides were separated using the Evosep One system (Evosep, Odense, Denmark) equipped with a 15 cm x 150 Dm i.d. packed with a 1.5 Dm Reprosil-Pur C18 bead column (Evosep Performance, EV-1137, PepSep, Marslev, Denmark) heated at 45°C with a butterfly sleeve oven (Phoenix S&T, Philadelphia, USA). The samples were run with a pre-programmed proprietary Evosep gradient of 44 min (30 samples per day) using water and 0.1% formic acid and solvent B acetonitrile and 0.1% formic acid as solvents. The LC was coupled to an Orbitrap Exploris 480 (Thermo Fisher Scientific, Bremen, Germany) using PepSep Sprayers and a Proxeon nanospray source. The peptides were introduced into the mass spectrometer via a PepSep Emitter 360 Dm outer diameter x 20 Dm inner diameter, heated at 300°C, and a spray voltage of 2 kV was applied. The injection capillary temperature was set at 300°C. The radio frequency ion funnel was set to 30%. For DIA data acquisition, full scan mass spectrometry (MS) spectra with a mass range of 350–1650 m/z were acquired in profile mode in the Orbitrap with a resolution of 120 000 FWHM. The default charge state was set to 2+, and the filling time was set at a maximum of 20 ms with a limitation of 3 x 10^6^ ions. DIA scans were acquired with 50 mass window segments of differing widths across the MS1 mass range. Higher collisional dissociation fragmentation (normalized collision energy 30%) was applied, and MS/MS spectra were acquired with a resolution of 30 000 FWHM with a fixed first mass of 200 m/z after accumulation of 1 x 10^6^ ions or after filling time of 45 ms (whichever occurred first). Data were acquired in profile mode. For data acquisition and processing of the raw data, Xcalibur 4.4 (Thermo) and Tune version 4.0 were used.

### Proteomic data processing

DIA raw data were analysed using the directDIA pipeline in Spectronaut (v.18 and v.19, Biognosys, Switzerland). Raw files were searched against a species-specific database (Homo sapiens, SwissProt release 2016_01) and a list of common contaminants (247 contaminants, including BSA, Trypsin, human keratins and other), using the default settings. For quantification, the default BGS factory settings are used, except for the following settings: proteotypicity filter = only protein group specific; major group quantity = median peptide quantity; major group top N = OFF; minor group quantity = median precursor quantity; minor group top N = OFF; data filtering = Q-value percentile with fraction = 0.2 and imputing strategy = global imputing; normalization strategy = global normalization; row selection = identified in all runs (complete). Differential abundance testing was performed in Spectronaut using a paired t-test between replicates. The data (candidate table) and data reports (protein quantities) were exported, and further data analyses and visualization were performed with R v.3.6 and RStudio server 2024.04.0 using in-house pipelines and scripts.

Zinc-binding proteins (Supplementary Table 5) are a list of manually curated proteins (in-house generated).

### Confocal imaging of labile zinc pools

For lysosome zinc imaging, cells were seeded in a four-chamber 35mm glass-bottom dish (Cellvis) and loaded with 200 nM FluoZin-3 AM and 75 nM LysoTracker Red DND-99 for 60 minutes in Hank’s balanced salt solution (HBSS). For mitochondria zinc imaging 5 DM Zinpyr-1 and 150 nM Mitotracker red or 0.1 Dmol/L MitoBright Deep red were loaded onto cells for 20 minutes in HBSS. Before imaging, the cells were washed with warmed PBS and DMEM with no phenol red, 10% FBS, and penicillin/streptomycin were added to remove fluorescence signal from phenol red. The dish containing the seeded cells was placed on a heated microscope stage in 5% CO_2_ and allowed to equilibrate for 15 minutes before the start of the experiment.

### Generation of KO lines using CRISPR-Cas9

Knockout cell lines were generated using CRISPR-Cas9 technology. ATG9A targeting sgRNA: 5’– GCAGCTGCTAGCCTCGCCCA-3’. ATG7 targeting sgRNA: 5’– GAAGCTGAACGAGTATCGGC-3’. Respective sgRNAs were cloned into the pX458 vector using GFP expression as a selectable marker. pX458 vectors were transfected into cells using PEI transfection reagent. 48 hrs post transfection GFP expressing cells were sorted into 96 well plates containing DMEM, 30% FBS, and penicillin/streptomycin. Plates were incubated for 4 weeks at 37°C in 5% CO_2_ and regularly monitored for colony formation. Colonies were expanded and knockout of ATG9A and ATG7 were validated using immunoblotting (ATG9A, Cell Signaling, 13509S), (ATG7, Cell Signaling, 8558S).

LONP1 knockout cells were generated using EditCo Gene Knockout kit using mixed guide sgRNAs. Mixed sgRNAs targeting LONP1: 5’-CGGGCUAUGGCGGCGAGCAC-3’, 5’– CGUCGCAGGUCCGCUGGCCU-3’, 5’-UGCUGGCCGCCGCCGGGGGG-3’. To form RNP complexes 180 pmol sgRNA and 20 pmol Cas9 were mixed with Nucleofection Reagent according to manufacturer’s specifications. PaTu-8988T cells were washed with PBS, dissociated and counted to determine cell density. 150,000 cells were combined with the newly formed RNP complexes and transferred to a Lonza Nucleocuvette and electroporated using the PANC1 electroporation program. After electroporation, cells were placed in a growth medium and transferred to a 24-well plate. The sgRNA target sites were amplified by PCR and sequenced. The genetic knockouts were confirmed using Synthego ICE CRISPR analysis, an indel deconvolution web-based tool. Additionally, we validated using immunoblotting (LONP1, Proteintech, 15440-1-AP).

### Co-Immunoprecipitation (Co-IP) of MT1-3xFLAG

PaTu-8988T cells stably expressing MT1-3xFlag were seeded onto 6-well plates at 1.0×10^6^ cells. The following day cells were depleted of zinc using 500 DM ZX1 for four hours after which the cells were harvested. The cells were washed with 1x PBS and then lysed with 160 Dl of RIPA buffer (25 mM Tris, 150 mM Sodium Chloride, 1% NP-40, 1% Sodium Deoxycholate. 0.1% SDS, pH 7.6) with protease inhibitor cocktail. Cell lysate was centrifuged for 10 min at max speed. The supernatant was collected into a new Eppendorf tube for Anti-Flag immunoprecipitation. 30 DL of the supernatant was kept as input. The remaining supernatant was diluted to 500 DL with 370 DL of RIPA buffer and added to 50 Dl of pre-washed anti-Flag Magnetic Agarose beads (Thermo Fisher Scientific). The lysates were incubated with the anti-Flag beads for 2 hours at 4°C with rotation. Following incubation, the beads were washed 3x with 1x PBS supplemented with a protease inhibitor cocktail. After the final wash beads were transferred to a new Eppendorf tube and remaining PBS was aspirated off the beads. Proteins were eluted from the beads using 100 DL of IgG elution buffer, pH 2.8 (Thermo Fisher Scientific). The beads were then incubated at 22°C and mixed using a 1000 rpm shaker for 10 minutes. The beads were put on a magnet for 30 seconds to separate magnetic beads from the supernatant. Eluted proteins in supernatant were then transferred to a new Eppendorf tube and 15 DL of 1M Tris pH 8.5 was used to neutralize acid pH of the elution buffer according to manufacturer’s instructions. Samples were flash frozen in liquid nitrogen and stored at –80°C for subsequent analysis.

### Seahorse Assay

Mitochondrial function and respiration were measured with the Agilent Seahorse XFe Mitochondria Stress test. The test was performed according to the manufacturer’s instructions. 8,000 cells were seeded in Seahorse XFe microplates and incubated at 37°C with 5% CO_2_ overnight. For the zinc-limited experiment, cells were treated with 500 DM ZX1 to chelate zinc ions in the culture media for the appropriate time. Cells were washed with Seahorse XF base assay medium and culture media was replaced with Seahorse medium supplemented with 20 mM glucose, 1 mM Pyruvate, and 2 mM L-glutamine. ZX1 was added to Seahorse medium in appropriate conditions to maintain zinc-limited conditions. Cells were incubated in a 0% CO_2_ incubator at 37°C for 1 hour to allow the medium to reach temperature and pH equilibrium before assay. Cells were treated with 1 DM oligomycin, 2 DM FCCP, and 0.5 DM rotenone. Oxygen Consumption Rate (OCR) was measured and analyzed using the Seahorse XFe report generator. Cells were then lysed, and total protein content was measured using Pierce BCA Assay. Total protein content was used to normalize OCR.

### Mouse treatment with zinc deficient diets

Experiments were performed using C57BL/6J wild-type mice from Jackson Laboratories. For routine colony maintenance, mice were fed a standard chow diet (Inotiv, Teklad Global 18% Protein Rodent Diet 2018). For LONP1 inhibition, mice were treated daily with 2.5 mg/kg CDDO-Me by intraperitoneal injection for three days before switching to zinc-free diet. To induce dietary zinc deficiency, mice were fed a zinc-free diet with 0.3 ppm zinc (Research Diets Inc., D25092402i) with a zinc-replete matched control diet of 29.5 ppm zinc (Research Diets Inc., D25092401i). Mice were fed zinc-free diet for six days and treated with 2.5 mg/kg CDDO-Me and euthanized. Liver tissue was collected in KPBS with protease and phosphatase inhibitors and homogenized.

## Data Analysis

Graphpad Prism 10 software was used for statistical analysis and data presentation. R was used for data presentation: R Packages: Tidyverse, ggplot2, ggrepel, readr, org.Hs.eg.db, enrichplot, DOSE, fastmap, Heatmap. Figures were assembled in Adobe Illustrator 2025. Sample size, error bars, and statistical tests are reported in the figure legends. Statistical tests utilized in this study include Fisher’s exact test, one-way analysis of variance (ANOVA), two-way ANOVA, and unpaired student’s t test or reported in the legend if different.

Key resource table.

| Key resource table |  |  |
| --- | --- | --- |
| Reagent | Source/Vendor | Identifier |
| LC3B | Cell Signaling | Cat# 2775S |
| SQSTM1/p62 | Cell Signaling | Cat# 5114S |
| Metallothionein antibody | Novus Biologics | Cat # NBP1-97493 |
| β-Actin | Santa Cruz Biotechnology | Cat# SC-47778 |
| ATG9A | Cell Signaling | Cat# 13509S |
| ATG7 | Cell Signaling | Cat# 8558S |
| Ubiquitin | Cell Signaling | Cat# 3936S |
| DYKDDDDK (Flag) tag | Cell Signaling | Cat# 14793S |
| LONP1 | Proteintech | Cat# 15440-1-AP |
| Anti-Mouse-HRP | Cell Signaling | Cat# 7076 |
| Anti-Rabbit HRP | Cell Signaling | Cat# 7077 |
| Bacteria strains |  |  |
| DH5alpha E.coli | Thermo Fisher Scientific | Cat# EC0112 |
| Chemicals, peptides, and recombinant proteins |  |  |
| CDDO-Me | Selleck Chemicals | Cat# S8078 |
| Protease inhibitor cocktail, in DMSO solution, For TC | Millipore Sigma | Cat# P1860 |
| Zinc replete Diet (29.5ppm) | Research Diets Inc | Cat# D25092401i |
| Zinc deficient diet (0.3ppm) | Research Diets Inc | Cat# D25092402i |
| DMEM, High Glucose | Fisher Scientific, Gibco | Cat# 11-965-118, 11965118 |
| Fetal bovine serum, (FBS, heat inactivated) | Thermo Fisher Scientific, Gibco | Cat# 10438026 |
| Penicillin/Streptomycin | Cytiva | Cat# SV30010 |
| Trypsin-EDTA | Gibco | Cat# 25300-062 |
| HBSS, no calcium, no magnesium | Fisher Scientific, Gibco | Cat# 14-175-095, 14175095 |
| Blasticidin | InvivoGen | Cat# ant-bl-1 |
| Puromycin dihydrochloride | Sigma-Aldrich | Cat# P7255 |
| XtremeGene9 DNA | Sigma-Aldrich | Cat#06365787001 |
| Transfection Reagent |  |  |
| PhosSTOP phosphatase inhibitor | Sigma-Aldrich | Cat# 04906837001 |
| PEI (polyethylenimine, MX 25kDa) | Polyscience | Cat# 23966-100 |
| Hoescht 33342 | Thermo Fisher Scientific | Cat# H3570 |
| Pierce™ RIPA Buffer | Thermo Fisher Scientific | Cat# 89900 |
| Seahorse XF Base Medium | Agilent Technologies | Cat# 103335-100 |
| Sodium Pyruvate (100mM) | Thermo Fisher Scientific | Cat# 11360070 |
| Pierce™ IgG Elution Buffer | Thermo Fisher Scientific | Cat# 21004 |
| Pierce™ Anti-DYKDDDDK Magnetic Agarose (Anti-Flag) | Thermo Fisher Scientific | Cat# A36797 |
| Lysotracker (red DND-99) | Thermo Fisher Scientific | Cat# L7528 |
| MitoBright LT Deep Red (MT12) | Dojindo | Cat# MT12-12 |
| Zinpyr-1 | Fisher Scientific | Cat# 50-596-113 |
| FluoZin-3 AM, cell Permeant | Thermo Fisher Scientific | Cat# F24195 |
| Mitoblock-6 | MedChem Express | Cat# HY-122652 |
| MitoTracker red FM | Thermo Fisher Scientific | Cat# M22425 |
| ZX1 (2-{[Bis(2-pyridinylmethyl)amino]ethylamino}benzenesulfonic acid hydrate sodium salt) | Strem Chemicals | Cat# 07-0350 |
| TPEN | Cayman Chemical Company | Cat #13340 |
| TETA (Triethylenetetramine dihydrochloride) | Millipore Sigma | Cat# T5033 |
| BCS (Bathocuproinedisulfonic acid disodium salt hydrate) | Sigma-Aldrich | Cat# 146625 |
| DFO (Deferoxamine (mesylate)) | Cayman Chemical Company | Cat# 14595 |
| Torin1 | Selleck Chemicals | Cat# S2827 |
| Bafilomycin A1 | Cayman Chemical Company | Cat# 11038 |
| Chloroquine Diphosphate | BioVision | Cat# 1825 |
| Bortezomib | Selleck Chemicals | Cat# S1013 |
| MG-132 | Selleck Chemicals | Cat# S2619 |
| FCCP | Selleck Chemicals | Cat# S8276 |
| BamHI-HF® Restriction Enzyme | New England Biolabs (NEB) | Cat# R3136S |
| NotI-HF® | New England Biolabs (NEB) | Cat# R3189S |
| XhoI | New England Biolabs (NEB) | Cat# R0146S |
| AgeI-HF | New England Biolabs (NEB) | Cat# R3552S |
| T4 DNA Ligase | New England Biolabs (NEB) | Cat# M0202T |
| Pierce™ Anti-HA Magnetic Beads | Thermo Fisher Scientific | Cat# 88837 |
| HEPES | Sigma-Aldrich | Cat# H3375 |
| Sodium chloride (NaCl) | Fisher Scientific, Millipore Sigma | Cat# 77-605, 77605 |
| EDTA | Sigma-Aldrich | Cat# 798681 |
| Triton™ X-100 | Sigma | Cat# X100 |
| Sodium Orthovanadate | MP Biomedicals | Cat# 0215966480 |
| Sodium Fluoride | Sigma-Aldrich | Cat# S1504 |
| Sodium Pyrophosphate | Sigma-Aldrich | Cat# 71515 |
| β-mercaptoethanol | Sigma-Aldrich | Cat# M3148 |
| Glycerol | Sigma-Aldrich | Cat# G5516 |
| Tris-Hcl | Sigma-Aldrich | Cat# PR-H5123 |
| Tris base | Fisher Scientific | Cat# BP1525 |
| Tween® 20 | Sigma-Aldrich | Cat# P9416 |
| Sodium Hydroxide | American bio | Cat# AB1916-01000 |
| CAPS | Sigma-Aldrich | Cat# C6070 |
| Ethanol | Gold Shield Distributors | Cat# 412811 |
| 16% Paraformaldehyde | VWR | Cat# 100503-917 |
| Saponin | Sigma-Aldrich | Cat# SAE0073 |
| Glycine | VWR | Cat# EM-4840 |
| Sodium dodecyl sulfate (SDS) | Sigma-Aldrich | Cat# 75746 |
| Immobilon-P PVDF Membrane | Sigma-Aldrich | Cat# IPVH00010 |
| Novex wedgewell Tris-Glycine Mini Gels | Thermo Fisher Scientific | Cat# XP04205BOX |
| Bovine Serum Albumin,heat shock fraction | Sigma-Aldrich | Cat# A7906 |
| Complete Protease Inhibitor Cocktail | Sigma-Aldrich | Cat# 05056489001 |
| Complete Protease Inhibitor Cocktail, mini | Sigma-Aldrich | Cat# 0215966480 |
| Pierce ECL 2 Western Blotting Substrate | Thermo Fisher Scientific | Cat# PI80196X3 |
| Urea | Bio-Rad | Cat# 161-0730 |
| Dithiothreitol (DTT) | Roth | Cat# 6908.3 |
| Iodoacetamide (IAA) | Sigma-Aldrich | Cat# I1149 |
| Phosphoric acid | AppliChem | Cat# A0989 |
| Triethylammonium biscarbonate buffer (TEAB) | Sigma-Aldrich | Cat# T7408 |
| Trypsin | Promega | Cat# V5111 |
| Formic acid | Biosolve | Cat# 0006914143BS |
| Acetonitrile | Biosolve | Cat# 0001204101BS |
| S-trap plate | Protifi | Cat# P002-96MCR |
| Evotips | Evosep | Cat# EV2013 |
| Isopropanol | Biosolve | Cat# 0016264101BS |
| Methanol | Biosolve | Cat# 0013684101BS |
| Commercially available kits |  |  |
| Mini-Prep Kit QIAprep spin | Qiagen | Cat# 27106 |
| Gel extraction Kit QIAEX II | Qiagen | Cat# 20021 |
| DNA Clean & Concentrator®-5 | Genesee Scientific | Cat# 11-302 |
| ZymoPURE II Plasmid Midiprep Kit | Genesee Scientific | Cat# 11-550B |
| Seahorse XFp FluxPak | Agilent Technologies | Cat# 103022-100 |
| SF Cell Line 4D-Nucleofector X Kit S | Lonza | Cat# V4XC-2032 |
| iRT peptides | Biognosys | Cat# Ki-3002-1 |

| Experimental models: Cell lines |  |  |
| --- | --- | --- |
| PaTu-8988T cells | ATCC | RRID:CVCL_1847 |
| U2OS cells | ATCC | HTB-96, RRID CVCL_0042 |

## Data Availability

All data used to evaluate the conclusions stated in the paper are presented in the paper and/or supplementary materials. Proteomics data have been deposited in the MassIVE repository and will be released upon peer review. Cell lines can be requested; these requests will be fulfilled by the corresponding author. Source data are provided with this paper

## Acknowledgements

We thank all members of the Abu-Remaileh Laboratory for their helpful discussions. We also thank Stanford University Cell Sciences Imaging Facility (CSIF). This work was supported by the National Institute of Health (NIH) Director’s New Innovator Award Program (1DP2CA271386), the Pew Charitable Trusts and the Alexander and Margaret Stewart Trust and the Knight Initiative for Brain Resilience at Stanford to M.A.-R. A.K.M. is supported by the Sarafan ChEM-H Chemistry/Biology Interface Program, O-Leary-Thiry Graduate Fellowship, and NIH T32 training grant (T32GM120007). The authors gratefully acknowledge support from the FLI Core Facilities Proteomics. A.O. is supported by the German Research Council (Deutsche Forschungsgemeinschaft, DFG) via the Research Training Group ProMoAge (GRK 2155) and the Chan Zuckerberg Initiative Neurodegeneration Challenge Network (award numbers 2020-221617, 2021-230967, and 2022-250618). A.O. and J.C.H. are supported by the Fritz-Thyssen Foundation (award number 10.20.1.022MN) and the NCL Stiftung. The FLI is a member of the Leibniz Association and is financially supported by the Federal Government of Germany and the State of Thuringia. J.C.H. is supported by the German Academic Exchange Service (DAAD) through its Postdoctoral Program. M.A.-R. is a Stanford Terman Fellow, a Sloan Fellow, and Pew-Steward Scholar for Cancer Research, supported by the Pew Charitable Trusts and the Alexander and Margaret Stewart Trust.

## Author Contributions

The study was conceptualized by A.K.M. and M.A.-R. A.K.M. and M.A.-R. developed the Methodology. A.K.M., J.C.H., A.O., H.T., M.J.M. conducted experiments. M.J.M. conducted mouse work. Data acquisition and analysis was conducted by A.K.M and J.C.H. Data analysis, data visualization and figure harmonization were performed by A.K.M.. A.K.M. and M.A.-R. wrote the original draft of this paper. All authors reviewed and edited the paper. M.A.-R. was responsible for funding acquisition

## Extended Data Figures

**Extended Data Figure 1.**
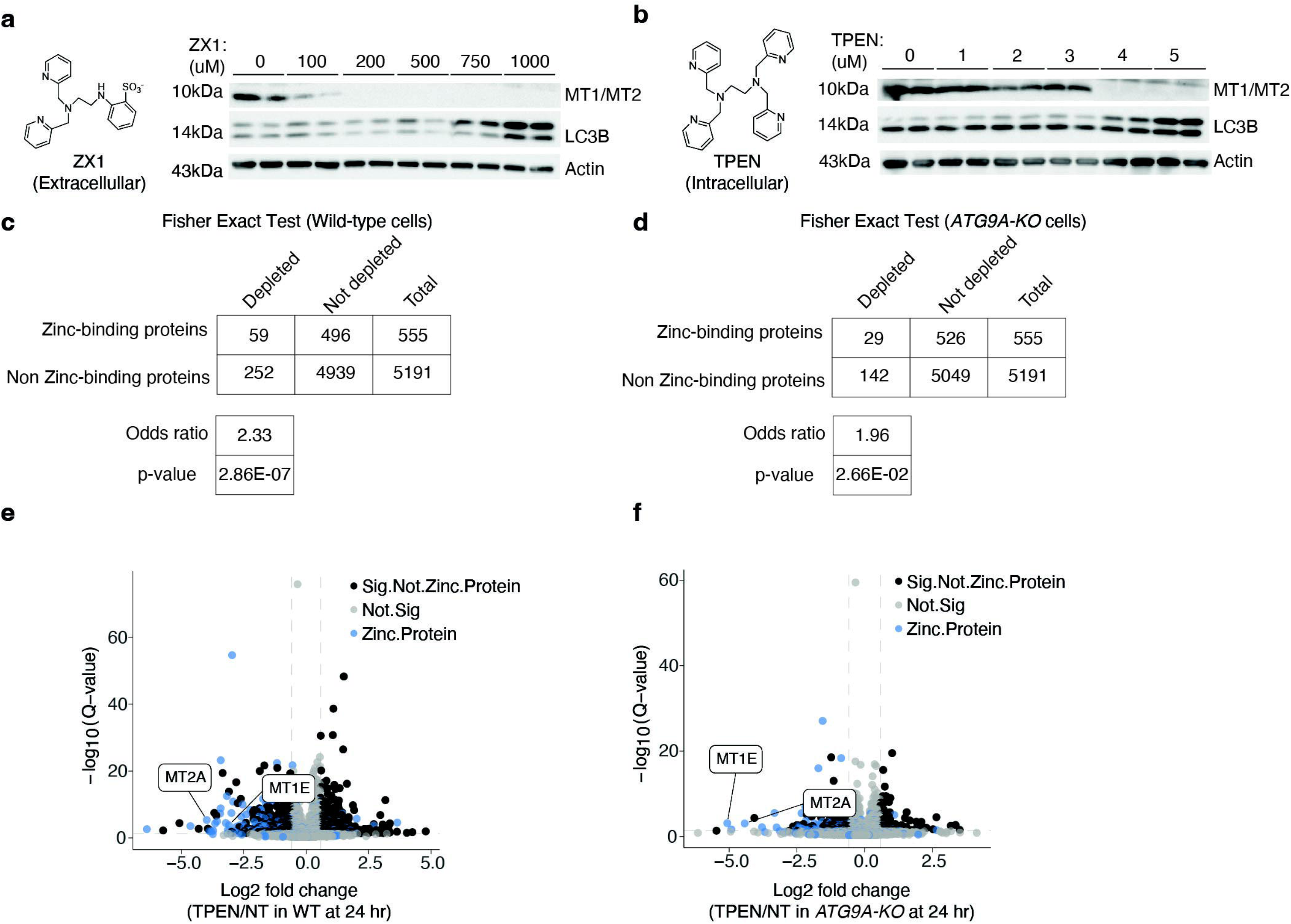
Optimization of Zinc limitation conditions. (a) Dose response of ZX1 treatment in wild-type PaTu-8988Ts for twenty-four hours. Left, chemical structure of ZX1. LC3B was used to monitor autophagy. Antibody recognizing MT1 and MT2 was used to monitor metallothioneins and D-Actin was used as loading control. (b) Dose response of TPEN treatment in wild-type PaTu-8988Ts for twenty-four hours. Left, chemical structure of TPEN. LC3B was used to monitor autophagy. Antibody recognizing MT1 and MT2 was used to monitor metallothioneins and D-Actin was used as loading control. (c) Fisher exact test showing ratio of zinc proteins depleted compared to total proteins depleted in twenty-four hour ZX1-treated wild-type PaTu-8988Ts. (d) Fisher exact test showing ratio of zinc proteins depleted compared to total proteins depleted in twenty-four ZX1-treated ATG9A-KO PaTu-8988Ts (e) Volcano plot of log2 fold changes (FC) in protein abundance in whole cell lysate of twenty-four hour TPEN treated wild-type (WT) PaTu-8988T cells compared to untreated condition (n = 3 biological replicates). Dashed lines indicate absolute log2 FC > 0.58 and Q < 0.05. Blue dots highlight zinc-binding proteins, black dots show other significantly affected proteins. (f) Volcano plot of log2 fold changes (FC) in protein abundance in whole cell lysate of twenty-four hour TPEN treated ATG9A-KO PaTu-8988T cells compared to untreated condition (n = 3 biological replicates). Dashed lines indicate absolute log2 FC > 0.58 and Q < 0.05. Blue dots highlight zinc-binding proteins, black dots show other significantly affected proteins.

**Extended Data Figure 2.**
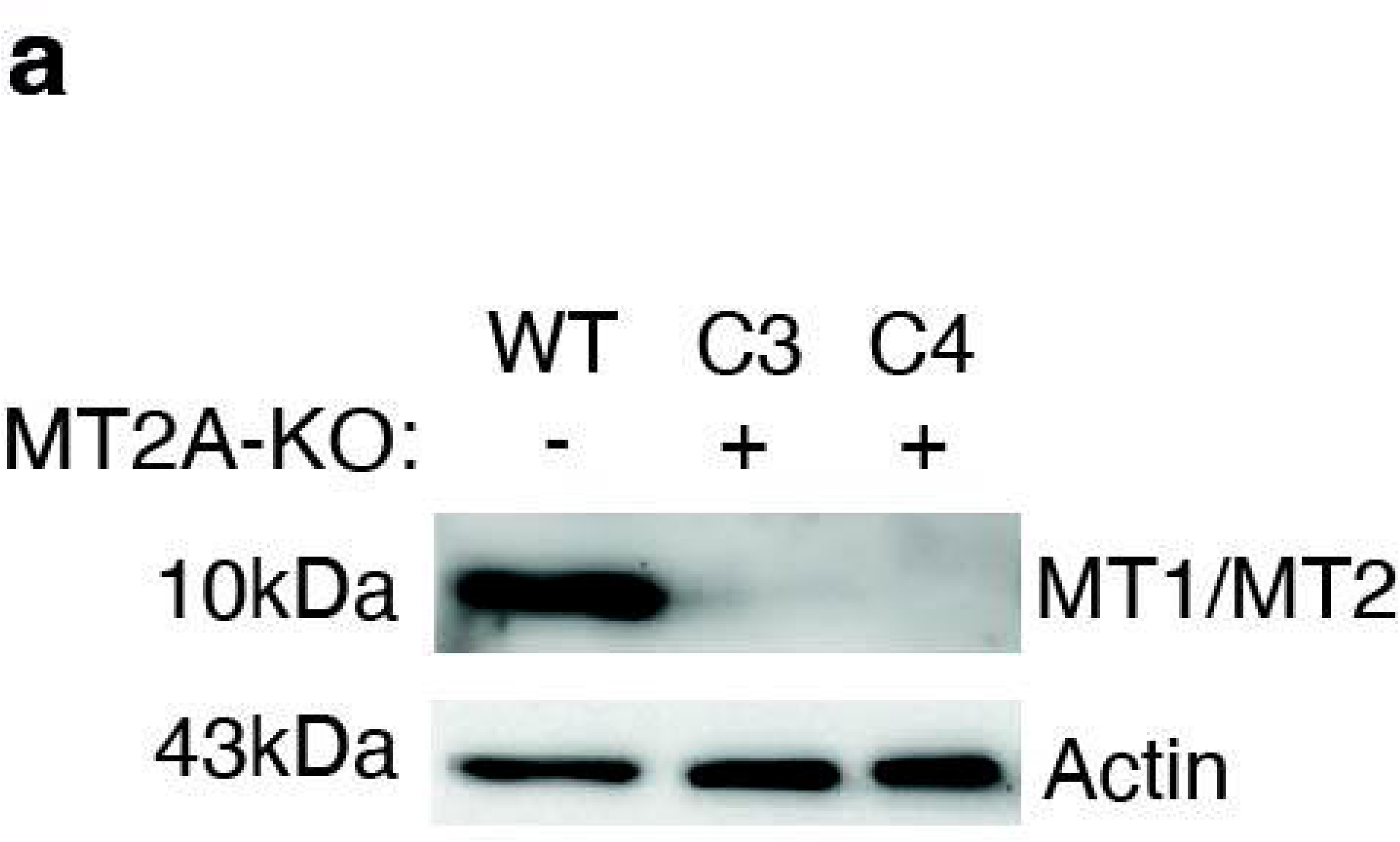
Validation of metallothionein antibody. (a) Immunoblot analysis to validate metallothionein antibody in wild-type and MT2A-KO PaTu-8988Ts. C3 and C4 are different KO clones. D-Actin was used as loading control.

**Extended Data Figure 3.**
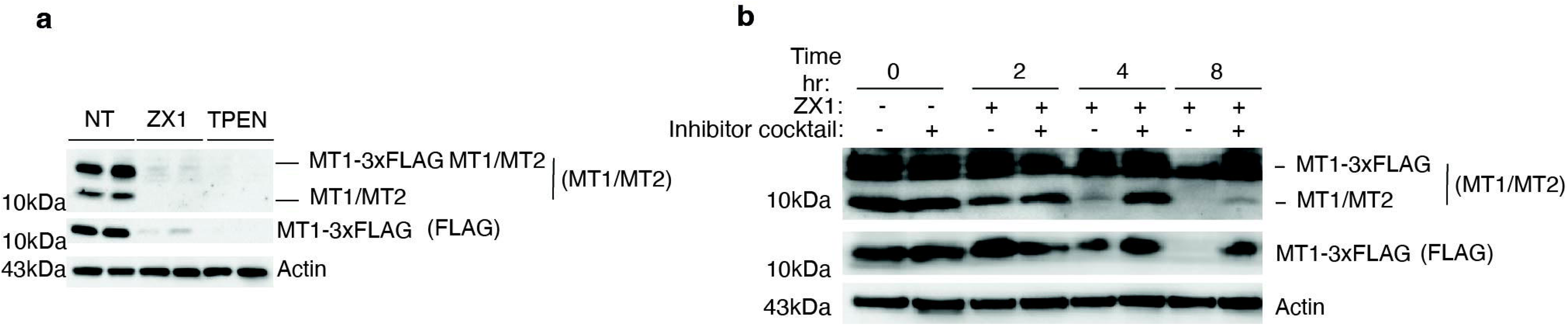
Validation of MT1-3xFLAG reporter response to zinc limitation. (a) Immunoblot analysis of MT1-3xFLAG expressing PaTu-8988T cells treated with zinc chelators ZX1 (500 DM) and TPEN (4 DM) at twenty-four hours. Antibody recognizing MT1 and MT2, as well as MT1-3xFLAG was used. MT1-3xFLAG was also monitored using anti-Flag antibody and D-Actin was used as loading control. (b) Immunoblot analysis of MT1-3xFLAG expressing PaTu-8988T cells treated with ZX1 in eight-hour time-resolved experiment and co-treated with commercial protease cocktail containing E-64, Leupeptin, Pepstatin A, Aprotinin, and Bestatin. Antibody recognizing MT1 and MT2, as well as MT1-3xFLAG was used. MT1-3xFLAG was also monitored using anti-Flag antibody and D-Actin was used as loading control.

**Extended Data Figure 4.**
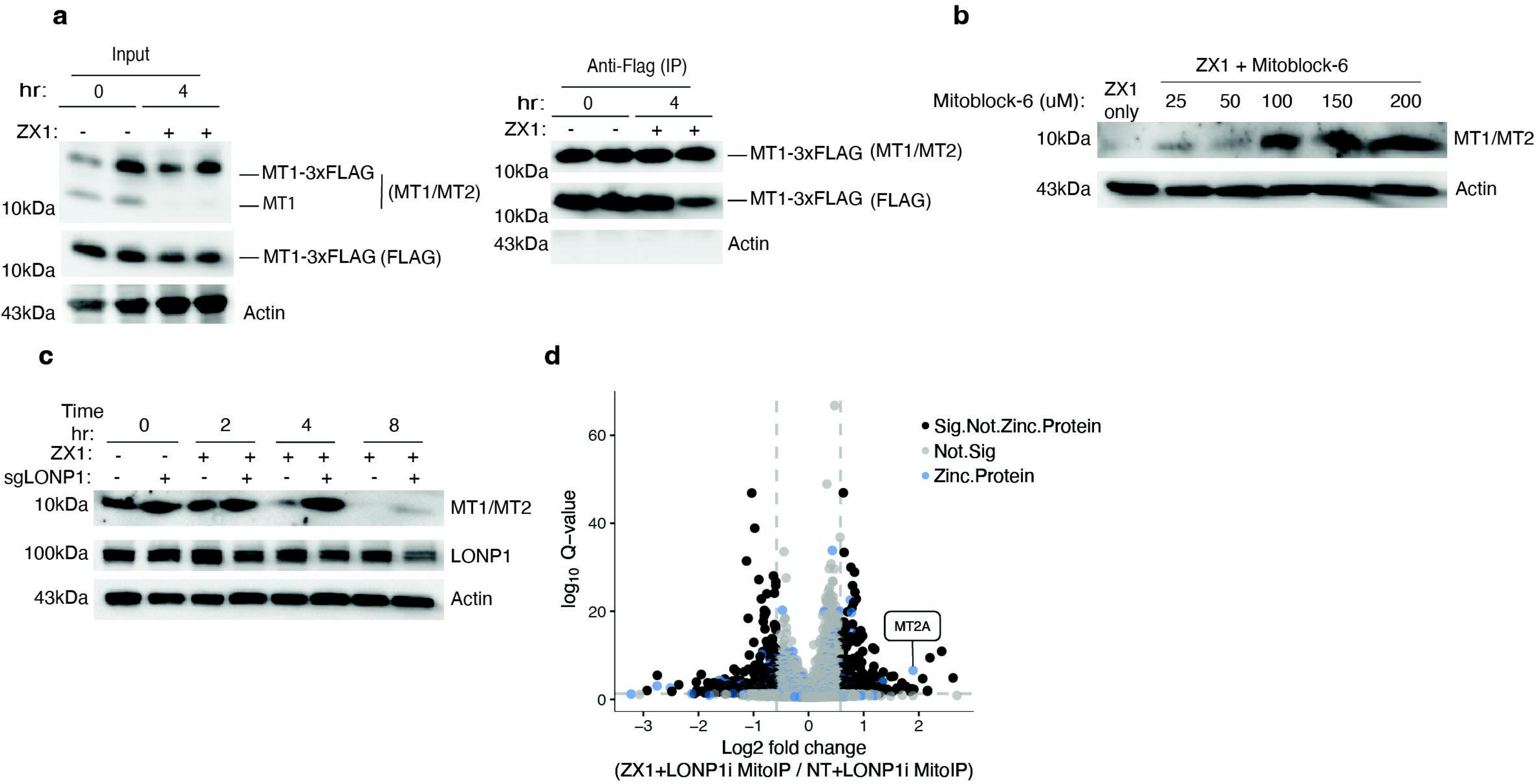
Metallothionein is imported into the mitochondria upon zinc starvation. (a) Immunoblot analysis of anti-Flag co-immunoprecipitation (CoIP) from PaTu-8988T cells expressing MT1-3xFLAG. Cells were treated with ZX1 (500 DM) for four hours. Input fraction on the left and IP fraction on the right. Antibody recognizing MT1 and MT2, as well as MT1-3xFLAG was used. MT1-3xFLAG was also monitored using anti-Flag antibody and D-Actin was used as loading control for the input fraction. (b) Immunoblot analysis of wild-type PaTu-8988T cells treated with ZX1 (500 DM) and the mitochondrial import blocker Mitoblock-6 (concentrations noted above) for eight hours. Antibody recognizing MT1 and MT2 was used and D-Actin was used as loading control. (c) Immunoblot analysis of wild-type PaTu-8988T cells as well as those electroporated with LONP1 sgRNA and were treated with ZX1 (500 DM) at the indicated time points. Antibodies recognizing MT1 and MT2 as well as LONP1 were used and D-Actin was used as loading control. (d) Volcano plot representation of log2 fold changes (FC) in protein abundance in mitoIP fraction from cells treated for four hours with ZX1 and co-treated with CDDO-Me (3 DM) compared to mitoIP from those in zinc replete media and co-treated with CDDO-Me (3 DM) (n = 4 biological replicates). Dashed lines indicate absolute log2 FC > 0.58 and Q < 0.05. Blue dots highlight zinc-binding proteins, black dots show other significantly enriched or depleted proteins.

## Supplementary Tables Legends

**Supplemental Table 1: Differential protein abundance of twenty-four hour ZX1 and/or TPEN treatment compared to no treatment (0hr)**

Table S1A: Differential protein abundance in wild-type PaTu-8988T cells treated with ZX1 for twenty-four hours versus no treatment

Table S1B: Differential protein abundance in ATG9A-KO PaTu-8988T cells treated with ZX1 for twenty-four hours versus no treatment

Table S1C: Differential protein abundance in wild-type PaTu-8988T cells treated with TPEN for twenty-four hours versus no treatment

Table S1D: Differential protein abundance in ATG9A-KO PaTu-8988T cells treated with TPEN for twenty-four hour versus no treatment

**Supplemental Table 2: Time course results of proteins whose abundance is significantly changing at twenty-four hours of ZX1 treatment in PaTu-8988T**

Table S1A: Differential protein abundance in wild-type PaTu-8988T cells treated with ZX1 at two, four, eight, sixteen, and twenty-four hours

Table S1B: Differential protein abundance in ATG9A-KO PaTu-8988T cells treated with ZX1 at two, four, eight, sixteen, and twenty-four hours

**Supplemental Table 3: Differential protein abundances of protein interactors of MT1-3xFLAG in Co-Immunoprecipitation (Co-IP)**

Table S3A: Differential protein abundances of protein interactors of MT1-3xFLAG at four hours of ZX1 treatment compared to PaTu-8988T without MT1-3xFLAG lysate background

**Supplemental Table 4: Mito-IP proteomics from ZX1-treated PaTu-8988T cells**

Table S4A: Differential protein abundances of Mito-IP from ZX1-treated PaTu-8988T cells that are co-treated with CDDO-Me compared to Mito-IP from cells treated only with CDDO-Me

**Supplemental Table 5: Manually curated list of zinc proteins**

Table S5A: List of zinc binding proteins that is manually curated

